# Single cell tools for WormBase

**DOI:** 10.1101/2021.07.04.451030

**Authors:** Eduardo da Veiga Beltrame, Valerio Arnaboldi, Paul W. Sternberg

**Affiliations:** Division of Biology and Biological Engineering, Caltech

## Abstract

We present two web apps for interactively performing common tasks with single cell RNA sequencing data: *scdefg* for interactive differential expression and *wormcells-viz* for visualization of gene expression. We made these tools available with public *C. elegans* datasets curated by WormBase at single-cell.wormbase.org. They can also be readily deployed for use with any other datasets using the source code available at github.com/WormBase/scdefg and at github.com/WormBase/wormcells-viz.

## Introduction

The number of single cell RNA sequencing (scRNA-seq) publications has exploded in recent years, with over 1200 studies currently available and over 350 new studies in 2020 alone [1]. This wealth of data is far from being used to it’s full potential, as most methods and tools for scRNA-seq data require programming proficiency in Python or R. Leveraging scRNA-seq data beyond the original studies requires tools that are easy to use and enable the integration, query, and display of data in ways that are useful for scientists. Over 85% of scRNA-seq studies use human or mouse samples, and the volume of data generated for these organisms is so high that their integration and unified management presents a formidable challenge by itself. But for other organisms such as *C. elegans*, for which there are only on the order of a dozen studies in the literature (Table 1), data curation, integration and maintenance of tools encompassing most of the published studies is manageable by a single individual or research group using simpler tools.

**Table 1:** Number of scRNA-seq studies for the most popular organisms. Data from [1] (www.nxn.se/single-cell-studies).

| Organism | Studies |
| --- | --- |
| Mouse | 514 |
| Human | 423 |
| Human & Mouse | 92 |
| Zebrafish | 25 |
| *Drosophila* | 12 |
| Rat | 10 |
| *C. elegans* | 6 |
| Yeast | 3 |
| *A. thaliana* | 3 |
| Chicken | 3 |
| *P. falciparum* | 3 |

In this spirit we present two tools: *scdefg*, for performing interactive differential expression (DE), and *wormcells-viz*, for visualization of gene expression data. These tools leverage the *anndata* file format [2], a standard file format for scRNA-seq data and annotated data matrices (anndata.readthedocs.io) and *scvi-tools* [3], a popular framework for generative modeling of scRNA-seq data and statistical analysis (scvi-tools.org).

## Overview of the *scdefg* and *wormcells-viz* tools

The *scdefg* app provides a single web page with an interface for performing DE on two groups of cells that can be selected according to the existing annotations in the data. For example, the user can select a group according to a combination of cell type, sample, tissue and experimental group. DE is computed using the scVI model [4] from scvi-tools [3], which enables quick computation even when using only CPUs. The results are displayed in the form of an interactive volcano plot (log fold change vs p-value) and MA plot (log fold change vs mean expression) that display gene descriptions upon mouseover, and sortable tabular results that can be downloaded in csv and Excel format. The app is written in Python using Flask and Plotly, and can be launched from the command line by specifying the path to a trained scVI model, plus the data labels by which cell groups may be stratified (e.g. cell type, experiment, sample). We have deployed the app on a cloud instance with only 8GB RAM and 2 vCPUs and observed this configuration is sufficient for handling a few concurrent users with results being returned in about 15 seconds.

The *wormcells-viz* app provides interactive and responsive visualizations of heatmaps, gene expression histograms and swarm plots. It is written in Javascript and Python and uses React (reactjs.org/) and D3 (d3js.org). Deploying the app requires having pre-computed gene expression values stored in three custom anndata files as described in the supplements. Using one anndata file for each visualization type keeps the codebase modular and simplifies adding more visualizations in the future. In the *wormcells-viz* documentation we provide a pipeline and tutorial to compute these expression values using scVI with any scRNA-seq dataset. The pipeline could be adapted for using other scRNA-seq analysis frameworks, but our recommendation is to use scvi-tools. The following visualizations are currently implemented.

### Heatmap

Visualization of mean gene expression in each group annotated in the data. The expression rates can be shown as either a traditional heatmap, or as a monochrome dotplot.

### Gene expression histogram

Histograms of the gene expression rates for a given gene across all cell types in the data. The histogram bin counts are computed from the scVI inferred expression rates for each cell.

### Swarm plot

This is a new visualization strategy to to facilitate candidate marker gene identification. For a given cell type, swarm plots visualize the relative expression of a set of genes across all cells annotated in a dataset. The Y axis displays the set of selected genes, and the X axis displays the log fold change in gene expression between the cell type of interest and all other cell types. This is computed by doing pairwise differential expression of each annotated cell type vs the cell type of interest.

**Figure 1:**
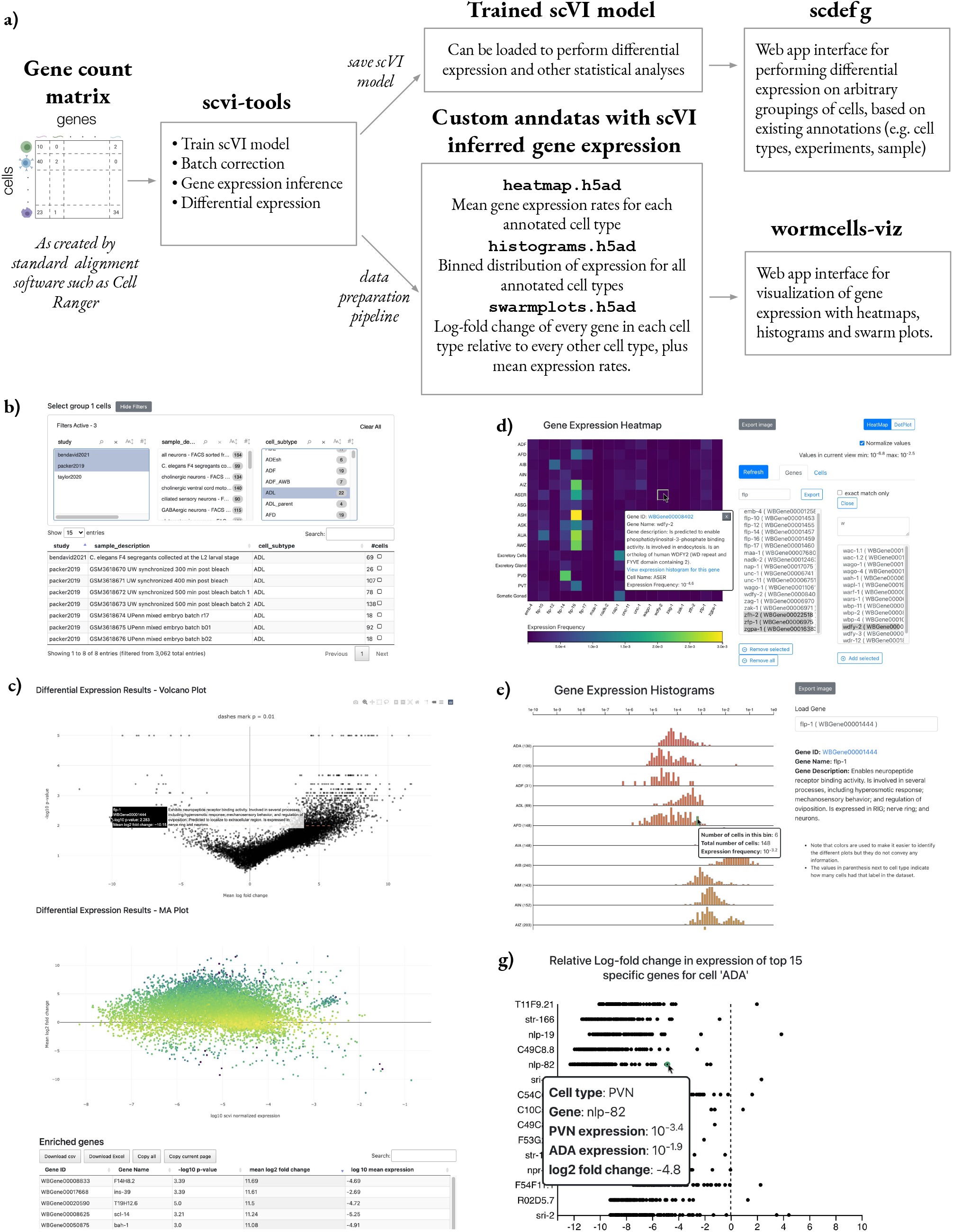
The WormBase single cell tools. **a)** Overview of the process to go from gene count matrix to deployment of the apps. Training a scVI model can be done quickly and only requires starting from the gene count matrix as outputted by standard alignment software such as Cell Ranger [5]. The *scdefg* app only requires as input the trained scVI model as saved by scvi-tools, while *wormcells-viz* requires using our pipeline to create the custom input anndatas, which are saved as. `h5ad` files (see supplements). **b)** The cell selection filter interface of *scdefg*. **c)** Results showing volcano plot, MA plot and part of the tabular results view of *scdefg*. **d)** Heatmap of *wormcells-viz*. **e)** Gene expression histogram of *wormcells-viz*, showing scVI normalized expression rates. **f)** Swarm plot of *wormcells-viz*.

### Rationale for using scvi-tools

The scvi-tools framework offers several models for single cell omics data, and for scRNA-seq in particular offers the scVI model [4], which is bayesian hierachical generative model that leverages variational autoencoders to enable robust statistical analysis. It is built with PyTorch (pytorch.org) and has been extensively validated [3]. Training the scVI model only requires a gene count matrix, as outputted by scRNA-seq alignment software such as Cell Ranger [5]. There are currently hundreds of software tools and pipelines developed for scRNA-seq data [6], and below we outline the considerations that led to our choice of using scvi-tools.

### Scalability

scvi-tools models readily scale datasets with millions of cells. Using a GPU even large models for millions of cells can be trained in a few hours, and models for a few thousand cells are trained in minutes. New datasets can also be integrated to an existing model without re-training from scratch.

### Consistent Development and Contributors

The scvi-tools codebase (github.com/YosefLab/scvi-tools) was first introduced in 2017, and published in 2018 [4], with consistent updates and improvements since [3]. It now boasts a mature and professional API and codebase that follows industry best practices, and has over 37 unique contributors and 56 releases.

### Extensible Framework for Analysis

Because the generative model of the data can be modified to reflect to capture our assumptions about underlying processes, the framework can be extended to model other aspects of scRNA-seq data. Currently, extensions include cell type classification and label transfer across batches, modelling single cell protein measurements, single cell chromatin accessibility assays, gene imputation in spatial data, and using a linear decoder to allow for interpretation of the learned latent space. Several peer reviewed articles have been published describing these extensions (see scvi-tools.org/press).

### WormBase deployments

At the moment, the majority of scRNA-seq studies generate data using the 10X Genomics Chromium technology [1]. This is also true for *C. elegans* scRNA-seq data. For the time being WormBase will focus development efforts on scRNA-seq tools on 10X Genomics data. Two considerations drive this:

i. Data integration of different batches with scvi-tools is more robust when there is more data, and when the technology and biological system of each batch is the same or similar. Attempting to integrate a small number of cells from unique technologies and unique biological systems can make it impossible to discern biological differences from technical artifacts.
ii. 10X Genomics supports the Cell Ranger [5] software pipeline for going from FASTQ files to gene count matrices. Technology standardization enables WormBase to uniformly reprocess FASTQ files in a single pipeline in the future.

### WormBase standard anndatas

We developed simple data wrangling guidelines for structuring anndata files with standard named fields, to be used by WormBase for scRNA-seq data curation, and enables straightforward usage in software pipelines. Following these guidelines, we curated high throughput scRNA-seq *C. elegans* datasets and made it available at Caltech Data (data.caltech.edu). See tables S1, S2 and S3 for details.

### Prospects for the Alliance of Genome Resources

WormBase is a member of the Alliance of Genome Resources (alliancegenome.org), a consortium of model organism databases that encompasses zebrafish, *Drosophila melanogaster*, mice, rat and yeast. In this work we have curated all of the available *C. elegans* data and made it available for the community. For other model organism databases it is also feasible to manually curate all the relevant public datasets. Once the data is curated, integrating such massive aggregated datasets with scvi-tools becomes straightforward. By leveraging the tools presented here it is possible to offer users an interface to query and compare data from several studies in a way that is quick and useful but without the need to write code.

### Code and data availability

- Web deployments with *C. elegans* scRNA-seq data: single-cell.wormbase.org
- Source code for *scdefg* and deployment instructions: github.com/WormBase/scdefg
- Source code for *wormcells-viz* and deployment instructions: github.com/WormBase/wormcells-viz.
- WormBase convention for anndata standard field names: github.com/WormBase/anndata-wrangling

### Providing user feedback

We want these tools and code to be useful to the community and encourage users to provide feedback and contributions. The authors may be contacted via email, WormBase users may submit a ticket to the WormBase helpdesk at wormbase.org/tools/support, and code contributions and suggestions can be made via pull requests or issues directly in the GitHub repositories.

## Acknowledgements

The authors would like to thank Paulo Nuin for reviewing the code, Tom Morrell for assistance with CaltechData, and Valentine Svensson for suggestions on the manuscript. This work was supported by grants NIH U24HG010859 and NIH U24HG002223.

## Supplementary Material

### Description of WormBase standard anndata

The Anndata file format (extension type .`h5ad`) was published in 2018 [2] as a generic class for handling annotated data matrices, with a focus on scRNA-seq data and Python support for machine learning, and integration with the SCANPY analysis frame-work (scanpy.readthedocs.io). Anndata is an efficient storage format because it uses HDF5 compression, and has come to be the standard format for manipulating scRNA-seq data in Python, as well as providing support in R (see github.com/theislab/zellkonverter and cran.r-project.org/web/packages/anndata/index.html).

Anndata’s popularity and uses continues to grow, with many packages standardizing their data maniulation around it. Examples include scvi-tools (scvi-tools.org), the Chan Zuckerberg cellxgene platform (chanzuckerberg.github.io/cellxgene) and the COVID-19 cell atlas initiative which standardized data distribution around anndata (covid19cellatlas.org).

Owing to the advantages of anndata and its popularity, WormBase adopted simple data wrangling guidelines for structuring published scRNA-seq data into anndata files with standard field names, in order to streamline their reuse in code pipelines. The WormBase anndata wrangling guidelines are described in tables S1 and S2 and maintained at github.com/WormBase/anndata-wrangling.

### Description of *wormcells-viz* custom anndatas

In this section we briefly review the conventions of the format expected by *wormcells-viz* for an anndata object instantiated in Python as `adata = anndata.AnnData`. A Jupyter notebook tutorial, and a Python script that goes from gene count matrices to the anndata files are described in the repository’s documentation at github.com/WormBase/wormcells-viz

### Gene expression histograms anndata formatting

These are used for plotting histograms of the scVI inferred expression rates for a given gene across all cell types in the data. Each anndata layer stores the expression values for all cell types and a given gene. The histogram bin counts are computed from the scVI normalized expression values, which are generated by a trained scVI model using the method `scvi.model.SCVI.get_normalized_expression`. The anndata `obs` contains the cell types and `var` contains the histogram bins, the genes are stored in layers with the keys being the gene ID. We store the genes in the layers because each view in the *wormcells-viz* app show the histograms for a single gene, so this makes accessing the data simpler. The histogram bin counts are computed from the log_10_ of the scVI expression rate. Each bin contains the number of cells in the dataset that were inferred to have an expression rate in the bin interval. There should be 100 bins with values between (−10, 0), representing expression rates from 10^−10^ to 10^0^. The data array is of shape *n*_*celltypes*_ × *n*_*bins*_ × *n*_*genes*_. The `adata` properties should be:

- `adata.obs` := Dataframe with cell types in index.
- `adata.var` := Dataframe with the bin intervals in index.
- `adata.X` := Not used.
- `adata.layers[gene_id]` := Each layer key is a gene ID, and contains a matrix with the binned expression rate counts for all cell types.
- `adata.uns` [‘ `about`’] := String with dataset information.

### Heatmap anndata formatting

These are used for plotting a heatmap of the average expression rates in each cell type, for a given selection of cell types and genes. The input data is a matrix of cell types and gene expression rates that contains the log_10_ scVI expression rate values. Cell types are in `adata.obs` and genes in `adata.var`. The data array is of shape *n*_*celltypes*_ × *n*_*genes*_. The `adata` properties should be:

- `adata.obs` := Dataframe with cell types in index.
- `adata.var` := Dataframe with gene IDs in index.
- `adata.X` := Matrix with log_10_ values of scVI normalized expression for each cell type and each gene.
- `adata.uns` [‘ `about`’] := String with dataset information.

### Swarm plots anndata formatting

These are used for plotting relative expression of a set of genes across all cells annotated in the dataset. The Y axis displays the set of selected genes, and X axis displays the log fold change of expression of that gene on all cell types relative to the cell of interest. The fold change is computed by doing pairwise differential expression of each annotated cell type vs the cell type of interest. For a given cell type, genes can be sorted by p-value, log fold change or mean expression rate. This part of the data is an array of shape *n*_*celltypes*_ ×*n*_*genes*_ ×*n*_*celltypes*_. The cell types are repeated along two dimensions, because this data contains the results of pairwise DE comparisons among each cell type in the data.

Plus *n*_*celltypes*_ matrices shaped like *n*_*celltypes*_ × *n*_*genes*_, because each unstructured layer `adata.uns[celltype]` contains a dataframe with global differential expression results for that cell type.

Finally, the unstructured layer `anndata.uns` [‘ `heatmap`’] contains a matrix with log_10_ scVI expression rates heatmap data (same data as used for plotting the heatmap), with genes in the index and cell types in the columns. This data is used to display the expression of each cell type on mouseover. The a`data` object properties should be:

- `adata.obs` := Dataframe with cell types in index.
- `adata.var` := Dataframe with gene IDs in index.
- `adata.X` := Not used.
- `adata.layers[cell_type]` := Mean log fold change for a given cell type for all genes.
- `adata.uns[cell_type]` :=The differential expression tables of the corresponding cell type vs all other cells. This can be used for ordering the genes by p-value, log fold change, and expression rate.
- `adata.uns` [‘ `heatmap`’] := Dataframe with genes in index and cell types in columns containing the log10 of the scVI expression frequency for each cell type
- `adata.uns` [‘ `about`’] := String with dataset information.

**Table S1:** WormBase naming guidelines for the anndata `var` annotation names - `anndata.AnnData.var` https://anndata.readthedocs.io/en/latest/anndata.AnnData.var.html

| var name | Description | Example value | Optionality |
| --- | --- | --- | --- |
| **index** | WormBase gene ID | WBGene00010957 | Required |
| **gene_id** | WormBase gene ID | WBGene00010957 | Required |
| **gene_name** | WormBase gene name | nduo-6 | Required |
| **gene_description** | WormBase short gene description. | Predicted to have NADH dehydrogenase activity. | Optional |

**Table S2:**
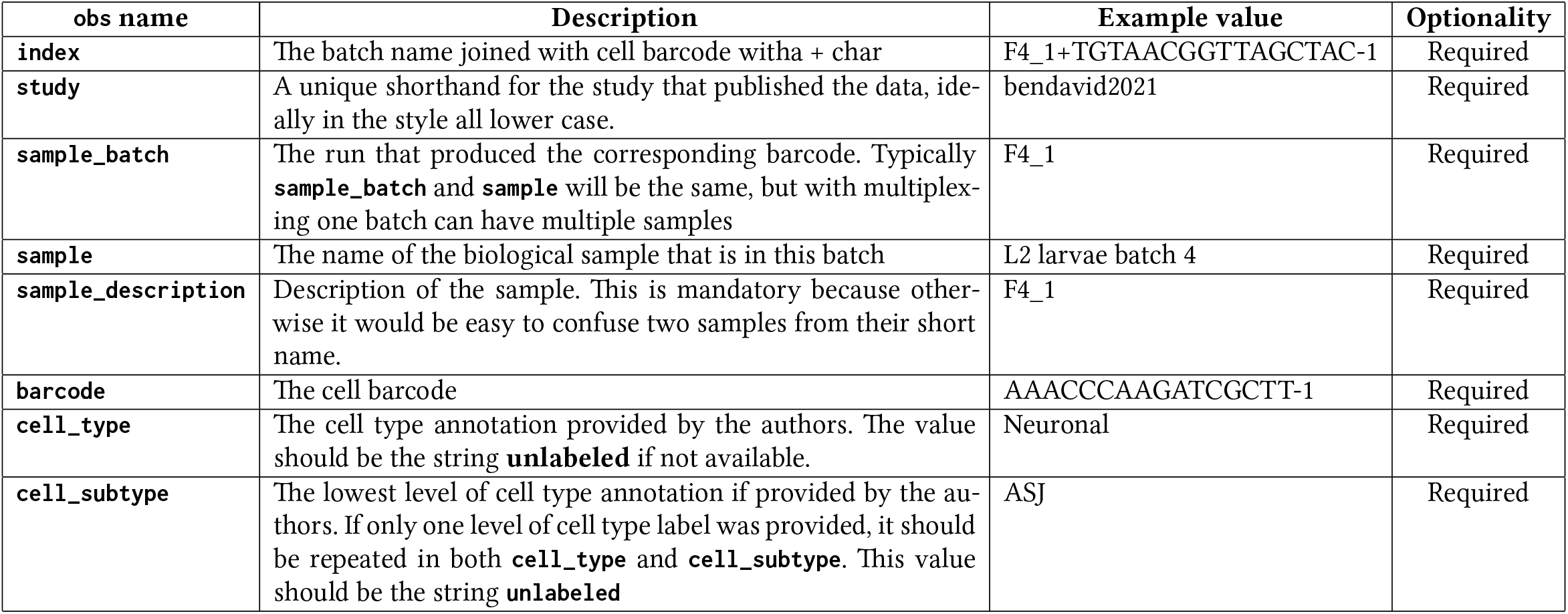
WormBase naming guidelines for the anndata `obs` annotation names - `anndata.AnnData.obs` https://anndata.readthedocs.io/en/latest/anndata.AnnData.obs.html

**Table S3:**
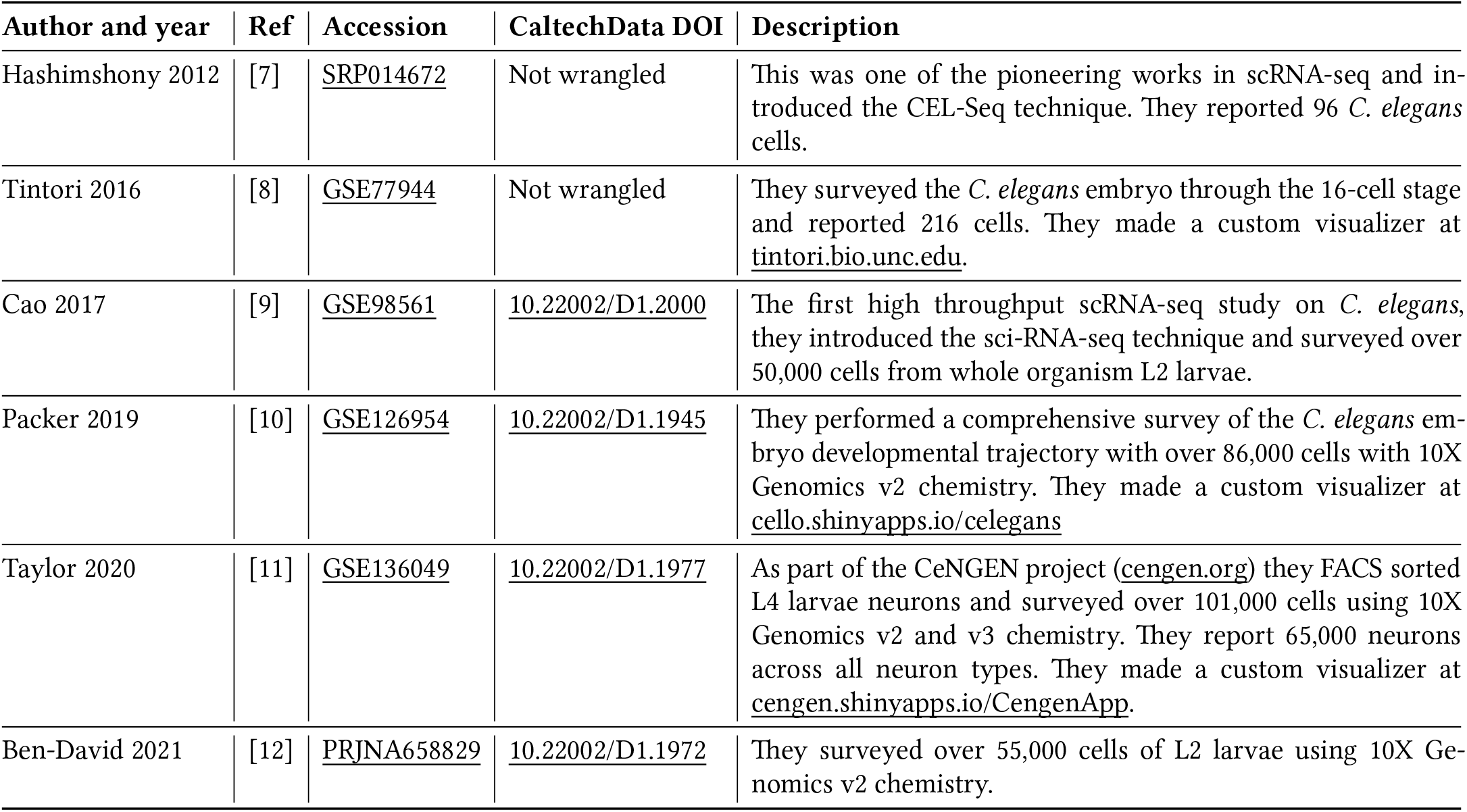
Summary of *C. elegans* single cell RNA sequencing datasets. High throughput data has been wrangled following the WormBase standard anndata convention and deposited at CaltechData (data.caltech.edu).

## References

[1] Svensson et al. (2020): A curated database reveals trends in single-cell transcriptomics, Database: 10.1093/database/baaa073.

[2] Wolf et al. (2018): SCANPY: large-scale single-cell gene expression data analysis, Genome Biology: 10.1186/s13059-017-1382-0.

[3] Gayoso et al. (2021): scvi-tools: a library for deep probabilistic analysis of single-cell omics data, bioRxiv: 10.1101/2021.04.28.441833.

[4] Lopez et al. (2018): Deep generative modeling for single-cell transcriptomics, Nature Methods: 10.1038/s41592-018-0229-2.

[5] Zheng et al. (2017): Massively parallel digital transcriptional profiling of single cells, Nat. Commun.: .

[6] Zappia et al. (2018): Exploring the single-cell RNA-seq analysis landscape with the scRNA-tools database, PLOS Comp. Biol.: 10.1371/journal.pcbi.1006245.

[7] Hashimshony et al. (2012): CEL-Seq: single-cell RNA-Seq by multiplexed linear amplification, Cell Reports: 10.1016/j.celrep.2012.08.003.

[8] Tintori et al. (2016): A Transcriptional Lineage of the Early C. elegans Embryo, Developmental Cell: 10.1038/10.1016/j.devcel.2016.07.025.

[9] Cao et al. (2017): Comprehensive single-cell transcriptional profiling of a multi-cellular organism, Science: 10.1126/science.aam8940.

[10] Packer et al. (2019): A lineage-resolved molecular atlas of C. elegans embryogenesis at single-cell resolution, Science: 10.1126/science.aax1971.

[11] Taylor, Seth R et al. (2020): Molecular topography of an entire nervous system, bioRxiv: 10.1101/2020.12.15.422897.

[12] Ben-David et al. (2021): Whole-organism eQTL mapping at cellular resolution with single-cell sequencing, Elife: 10.7554/eLife.65857.

